# Changes of urine proteome after intragastric administration of polysaccharide iron complex in rats

**DOI:** 10.1101/2024.03.05.583147

**Authors:** Ziyun Shen, Minhui Yang, Haitong Wang, Youhe Gao

## Abstract

Iron is an essential trace element to maintain the normal physiological function of organisms. In this study, the urine proteome of rats before and after short-term intragastric administration of polysaccharide-iron complex (28mg/kg/d iron, which is equivalent to the dose of anemia prevention in adults) was compared and analyzed by using two analysis methods: individual comparison and group comparison. Many different proteins were reported to be related to iron, including 2’, 3’ -cyclic nucleotide 3’ -phosphodiesterase (CNPase) (7.7 times higher than that after gavage, p=0.0039), p38 (14.5 times higher than that before gavage, p=0.003), etc. In the individual comparison, Hepcidin was up-regulated in 4 rats simultaneously. The biological processes of differential protein enrichment include carbohydrate metabolism, iron ion reaction, apoptosis regulation, hematopoietic progenitor cell differentiation, etc. Molecular functions (e.g., complement binding, hemoglobin binding, etc.), KEGG pathways (e.g., complement and coagulation cascade, cholesterol metabolism, malaria, etc.) have also been shown to be associated with iron. This study contributes to the in-depth understanding of the biological function of iron from the perspective of urine proteomics, and provides a new research perspective for the prevention, diagnosis, treatment and monitoring of iron-related disorders.

## 1 Introduction

Trace elements play an indispensable role in various physiological processes of the organism. In recent years, with the in-depth study of the role of trace elements, scientists have come to realize that the homeostasis of trace elements is also related to the pathogenesis of many diseases.

Iron is one of the trace elements necessary for maintaining normal physiological functions of living organisms, and it is involved in a variety of important biological processes in the body, such as oxygen transport, cellular respiration, and DNA synthesis. Disturbances in iron metabolism may lead to an imbalance in the biochemical balance of the organism, leading to a series of health problems^[1]^.

Since urine is not part of the internal environment, in contrast to plasma, there is no mechanism for homeostasis, and it is able to accumulate early changes in the physiological state of the organism, reflecting more sensitively the changes in the organism, and is a source of next-generation biomarkers^[2]^. The proteins in urine contain a wealth of information that can reflect the small changes produced in different systems and organs of the organism.

Our laboratory has previously reported that the urine proteome is able to reflect the effects of magnesium threonate intake on the body in a more systematic and comprehensive manner, with the potential to provide clues for clinical nutrition research and practice^[3]^. However, to date, there have been no studies exploring the effects of iron on the body from the perspective of the urine proteome.

In the present study, Polysaccharide-Iron Complex (PIC) was selected as an iron supplement, which can rapidly increase blood iron levels with elevated hemoglobin. It is mildly irritating to the gastrointestinal mucosa, has few adverse effects, can be administered continuously, has a high absorption rate, and is used for the prevention and treatment of iron deficiency anemia. The aim of this study was to investigate the changes in the urinary proteome of rats after short-term intake of polysaccharide-iron complexes to further understand the biological functions of iron in organisms and its overall effects, and to provide new research perspectives for nutritional studies.

## 2 Materials and Methods

### 2.1 Experimental materials

#### 2.1.1 Laboratory consumables

5 ml sterile syringe (BD), gavage needle (16-gauge, 80 mm, curved needle), 1.5 ml/2 ml centrifuge tube (Axygen, USA), 50 ml/15 ml centrifuge tube (Corning, USA), 96-well cell culture plate (Corning, USA), 10 kD filter (Pall, USA), Oasis HLB solid phase extraction column (Waters, USA), 1ml/200ul/20ul pipette tips (Axygen, USA), BCA kit (Thermo Fisher Scientific, USA), high pH reverse peptide separation kit (Thermo Fisher Scientific, USA), iRT ( indexed retention time, BioGnosis, UK).

#### 2.1.2 Experimental apparatus

Rat metabolic cages (Beijing Jiayuan Xingye Science and Technology Co., Ltd.), frozen high-speed centrifuge (Thermo Fisher Scientific, USA), vacuum concentrator (Thermo Fisher Scientific, USA), DK-S22 electric thermostatic water bath (Shanghai Jinghong Experimental Equipment Co., Ltd.), full-wavelength multifunctional enzyme labeling instrument (BMG Labtech, Germany), oscillator (Thermo Fisher Scientific, USA), TS100 constant temperature mixer (Hangzhou Ruicheng Instruments Co. BMG Labtech), electronic balance (METTLER TOLEDO, Switzerland), -80 □ ultralow-temperature freezing refrigerator (Thermo Fisher Scientific, USA), EASY-nLC1200 Ultra High Performance Liquid Chromatography system (Thermo Fisher Scientific, USA), and Orbitrap Fusion Lumos Tribird Mass Spectrometer (Thermo Fisher Scientific, USA) were used.

#### 2.1.3 Experimental reagents

Polysaccharide Iron Complex Capsules (Drug License H20030033) were manufactured by Shanghai Pharmaceutical Group Qingdao Guofeng Pharmaceutical Branch Co. In addition, Trypsin Golden (Promega, USA), dithiothreitol (DTT) (Sigma, Germany), iodoacetamide (IAA) (Sigma, Germany), ammonium bicarbonate NH4HCO3 (Sigma, Germany), urea (Sigma, Germany), and purified water (Wahaha, China) were used, Methanol for mass spectrometry (Thermo Fisher Scientific, USA), Acetonitrile for mass spectrometry (Thermo Fisher Scientific, USA), Purified water for mass spectrometry (Thermo Fisher Scientific, USA), Tris-Base (Promega, USA), Thiourea (Sigma, Germany) and other reagents.

#### 2.1.4 Analytical software

Proteome Discoverer (Version2.1, Thermo Fisher Scientific, USA), Spectronaut Pulsar (Biognosys, UK), Ingenuity Pathway Analysis (Qiagen, Germany); R studio (Version1.2.5001); Xftp 7; Xshell 7.

### 2.2 Experimental Methods

#### 2.2.1 Animal modeling

In this study, 17-week-old rats were used to minimize the effects of growth and development during gavage. Five healthy SD (Sprague Dawley) 9-week-old male rats (250±20 g) were purchased from Beijing Viton Lihua Laboratory Animal Technology Co. The rats were kept in a standard environment (room temperature (22±2)℃, humidity 65%-70%) for 8 weeks and weighed 500-600 g. The experiments were started, and all experimental operations followed the review and approval of the Ethics Committee of the School of Life Sciences, Beijing Normal University.

Dietary nutrient tolerable upper intake levels (UL): the average daily maximum intake of a nutrient for a population of a certain physiological stage and gender, which has no side effects or risks to the health of almost all individuals. Recommended nutrient intakes (RNI): intake levels that meet 97-98% of the individual needs of a particular age, gender, or physiological group.

According to the Dietary Guidelines for Chinese Residents, the recommended daily intake (RNI) of iron is 20mg/d, and the tolerable upper intake levels (UL) is 42mg/d^[4]^. In this study, the dose of iron in rats by gavage was 28mg/kg/d, which is equivalent to the dose for adults. equivalent to the dose used to prevent anemia in adults. A gavage solution was prepared by dissolving 3 g of polysaccharide iron complex (about 1.4 g of iron content) in 500 ml of sterile water. Each rat was gavaged with 5 ml of iron polysaccharide solution once a day for 4 days. The first day of gavage was recorded as Fe-D1, and so on. Sampling time points were set up before and after gavage for their own before and after control, the samples collected on the day before gavage were the control group, recorded as Fe-D0, sample number 51-55, the samples collected on the fourth day of gavage were the experimental group, recorded as Fe-D4 sample number 61-65.

#### 2.2.2 Urine sample collection

One day before the start of gavage of polysaccharide-iron complex (D0) and 4 days after the gavage of polysaccharide-iron complex (D4), each rat was individually placed in a metabolic cage at the same time, fasted and dehydrated for 12 h, and urine was collected overnight; urine samples were collected and placed in the refrigerator at -80°C for temporary storage.

#### 2.2.3 Urine sample processing

Two milliliters of urine sample was removed, thawed, and centrifuged at 4 °C and 12,000×g for 30 minutes. The supernatant was removed, and 1 M dithiothreitol (DTT, Sigma) storage solution (40 µl) was added to reach the working concentration of DTT (20 mM). The solution was mixed well and then heated in a metal bath at 37 °C for 60 minutes and allowed to cool to room temperature.

Then, iodoacetamide (IAA, Sigma) storage solution (100 µl) was added to reach the working concentration of IAM, mixed well and then reacted for 45 minutes at room temperature protected from light. At the end of the reaction, the samples were transferred to new centrifuge tubes, mixed thoroughly with three times the volume of precooled anhydrous ethanol, and placed in a freezer at -20 °C for 24 h to precipitate the proteins.

At the end of precipitation, the sample was centrifuged at 4 °C for 30 minutes at 10,000×g, the supernatant was discarded, the protein precipitate was dried, and 200 µl of 20 mM Tris solution was added to the protein precipitate to reconstitute it. After centrifugation, the supernatant was retained, and the protein concentration was determined by the Bradford method.

Using the filter-assisted sample preparation (FASP) method, urinary protein extracts were added to the filter membrane of a 10-kD ultrafiltration tube (Pall, Port Washington, NY, USA) and washed three times with 20 mM Tris solution. The protein was resolubilized by the addition of 30 mM Tris solution, and the protein was added in a proportional manner (urinary protein:trypsin = 50:1) to each sample. Trypsin (Trypsin Gold, Mass Spec Grade, Promega, Fitchburg, WI, USA) was used to digest proteins at 37 °C for 16 h.

The digested filtrate was the peptide mixture. The collected peptide mixture was desalted by an Oasis HLB solid phase extraction column, dried under vacuum, and stored at -80 °C. The peptide mixture was then extracted with a 0.1% peptide mixture. The lyophilized peptide powder was redissolved by adding 30 μL of 0.1% formic acid water, and then the peptide concentration was determined by using the BCA kit. The peptide concentration was diluted to 0.5 μg/μL, and 4 μL of each sample was removed as the mixed sample.

#### 2.2.4 LC-MS/MS analysis

All identification samples were added to a 100-fold dilution of iRT standard solution at a ratio of 20:1 sample:iRT, and the retention times were standardized. Data-independent acquisition (DIA) was performed on all samples, and each sample measurement was repeated 3 times, with 1-mix samples inserted after every 10 runs as a quality control. The 1-µg samples were separated using EASY-nLC1200 liquid chromatography (elution time: 90 min, gradient: mobile phase A: 0.1% formic acid, mobile phase B: 80% acetonitrile), the eluted peptides were entered into the Orbitrap Fusion Lumos Tribird mass spectrometer for analysis, and the corresponding raw files of the samples were generated.

#### 2.2.5 Data processing and analysis

The raw files collected in DIA mode were imported into Spectronaut software for analysis, and the highly reliable protein standard was peptide q value<0.01. The peak area quantification method was applied to quantify the protein by applying the peak area of all fragmented ion peaks of secondary peptides, and the automatic normalization was processed.

Proteins containing two or more specific peptides were retained, and missing values were replaced with 0. The amount of different proteins identified in each sample was calculated, and samples from rats prior to gavage of the polysaccharide-iron complex were compared with samples from rats 4 days after gavage of the polysaccharide-iron complex to screen for differential proteins.

Unsupervised cluster analysis (HCA), principal component analysis (PCA), and OPLS-DA analysis were performed using the Wukong platform (https://omicsolution.org/wkomics/main/). Functional enrichment analysis of differential proteins was performed using the DAVID database (https://david.ncifcrf.gov/) to obtain results in 3 areas: biological process, cellular localization and molecular function. Differential proteins and related pathways were searched based on Pubmed database (https://pubmed.ncbi.nlm.nih.gov/). Protein interaction network analysis was performed using the STRING database (https://cn.string-db.org/).

#### 2.2.6 Randomized group analysis

When using histology techniques to study disease biomarkers, it is common to screen for differences between disease and control groups. Due to the large amount of histologic data and limited sample size, differences between the two groups may arise randomly. For this reason, we use a randomized grouping statistical analysis strategy, which is applicable to the study of disease biomarkers in clinical histology with limited sample sizes, and determine whether the differences between the two groups are randomly generated^[5]^.

A total of 10 samples before (n=5) and after (n=5) gavage were randomly divided into two groups, and the mean of the number of differential proteins in all randomized combinations was calculated according to the same screening conditions.

#### 2.2.7 Analysis of differential proteins and functional annotations using the Pubmed database

In Pubmed for differential proteins and functional annotations were searched and analyzed, the specific search condition was to include both the keyword and iron in the title or abstract, for example, “iron [Title/Abstract] AND heme [Title/Abstract]”. These articles were then read and screened one by one to analyze the association of differential proteins, as well as molecular functions, biological processes, and pathways enriched by differential proteins, with iron.

## 3 Results and discussion

### 3.1 Characteristics of rats after gavage of polysaccharide-iron complexes

In the course of the experiment, we observed the drinking, feeding, body weight, hair and other characteristics of rats after gavage of polysaccharide iron complex. It was found that before and after gavage of the polysaccharide iron complex, the body weight of the rats remained basically stable, and their water intake, food intake and activities were normal. After gavage of the polysaccharide-iron complex, the rats’ feces were dark in color, and their hair was messy, which might be caused by excessive iron intake.

### 3.2 Urine protein identification profile and unsupervised cluster analysis

A total of 1,803 proteins were identified in the urine samples from before the gavage of polysaccharide-iron complex (D0) and on day 4 of the gavage (D4) (satisfy Unique peptides>1, FDR<1%). Unsupervised cluster analysis (HCA) and principal component analysis (PCA) were performed on the total proteins, and the results are shown in Figs. 2 and 3. The results of HCA and PCA showed relatively significant changes in the urinary proteome of the rats after ingestion of the polysaccharide-iron complexes, which may reflect the rapid response of the organism to exogenous iron. However, the distribution of sample points was scattered, indicating some inter-individual differences.

**Figure 1.**
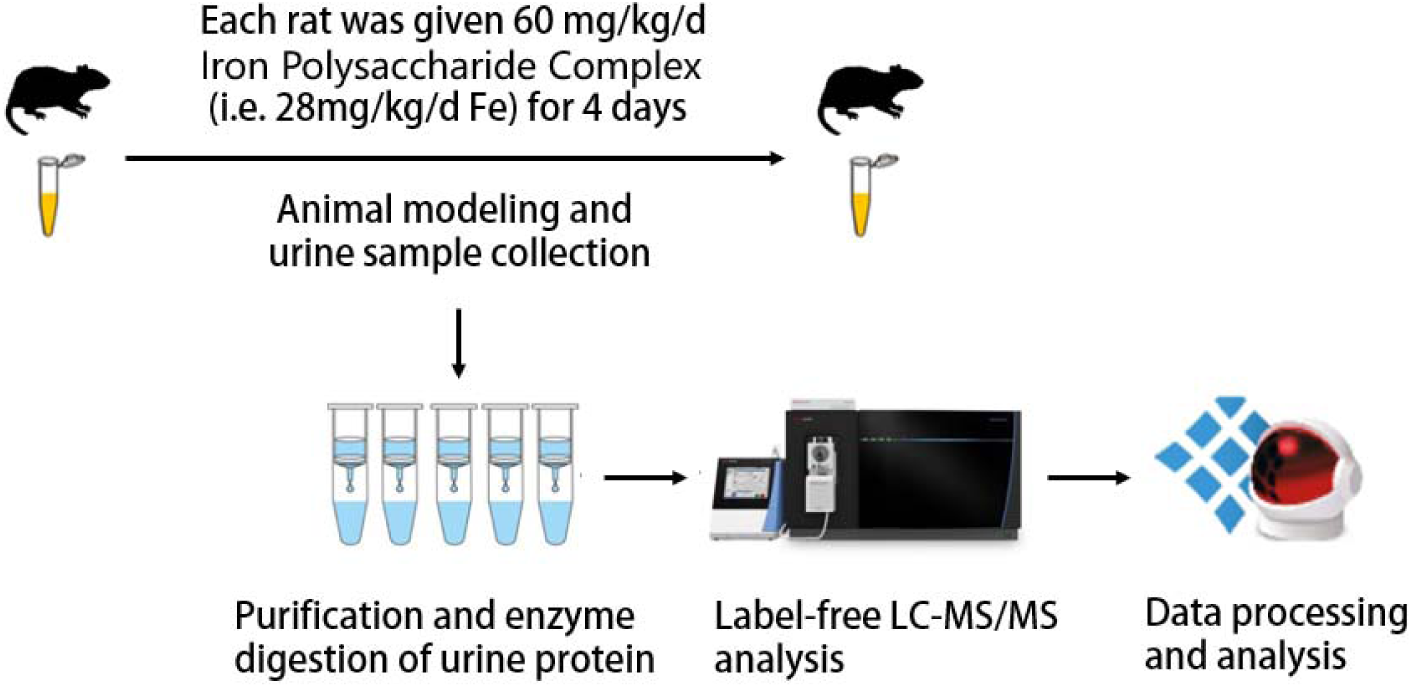
Research methodology and technical route

**Figure 2.**
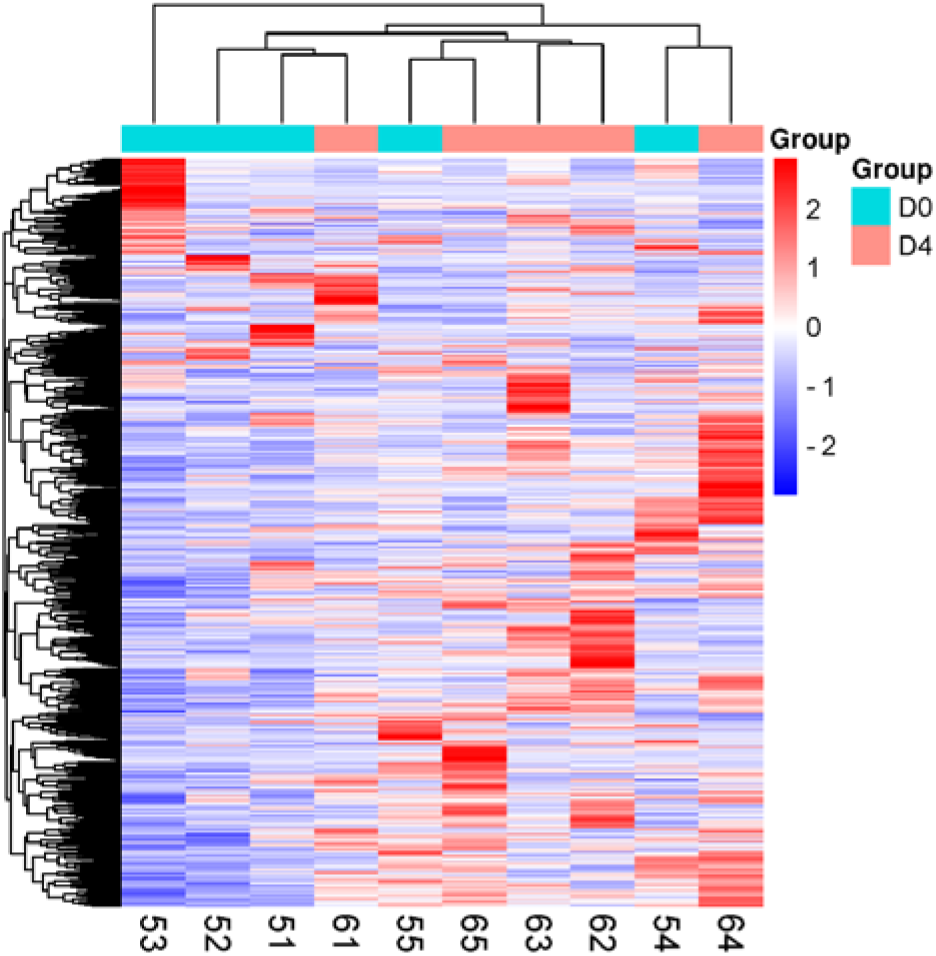
Unsupervised cluster analysis (HCA) of total protein in urine samples before gavage of polysaccharide-iron complexes versus day 4 of gavage

**Figure 3.**
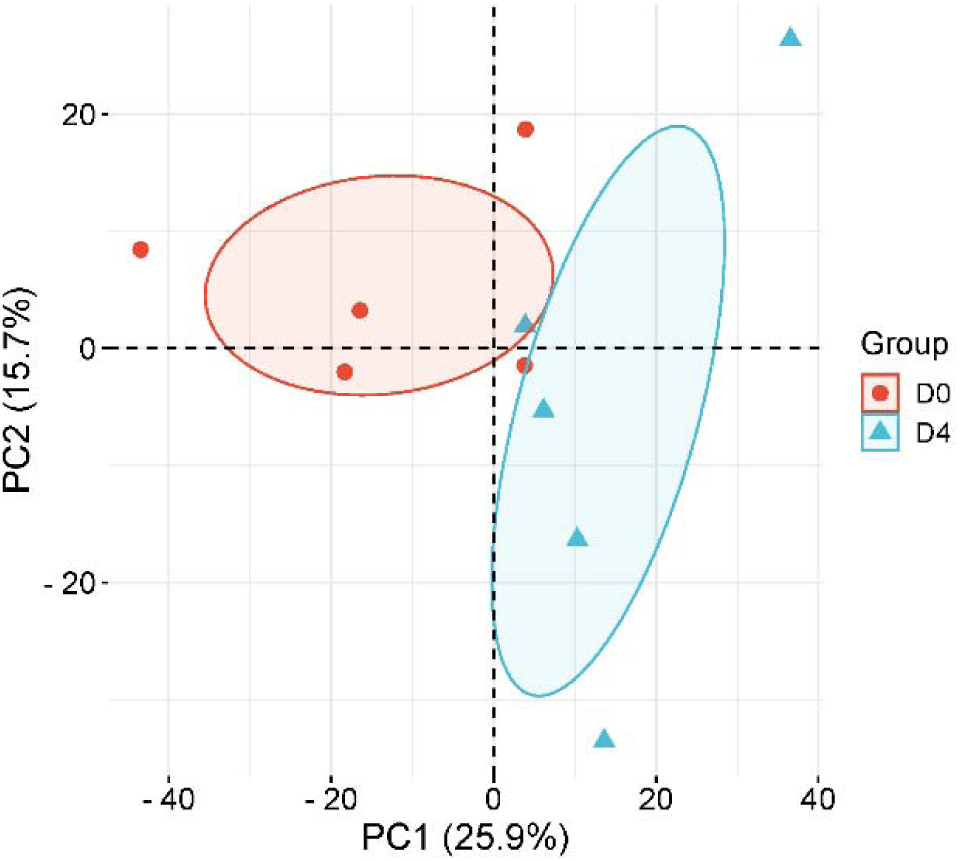
Principal component analysis (PCA) of total protein in urine samples before gavage of polysaccharide-iron complexes versus day 4 of gavage

### 3.3 Comparison of groups

#### 3.3.1 Differential protein analysis

Missing values were replaced with 0, and 157 differential proteins were screened by comparing the pre-gavage samples of rats with the samples on day 4 of gavage in groups. Screening conditions for differential proteins were: t-test analysis P value <0.05, Fold change (FC) >1.5 or <0.67. See Supplementary Table for details.

Among them, there were 52 differential proteins with P-value <0.01 and highly significant changes before and after gavage, as shown in Table 1.

**Table 1.**
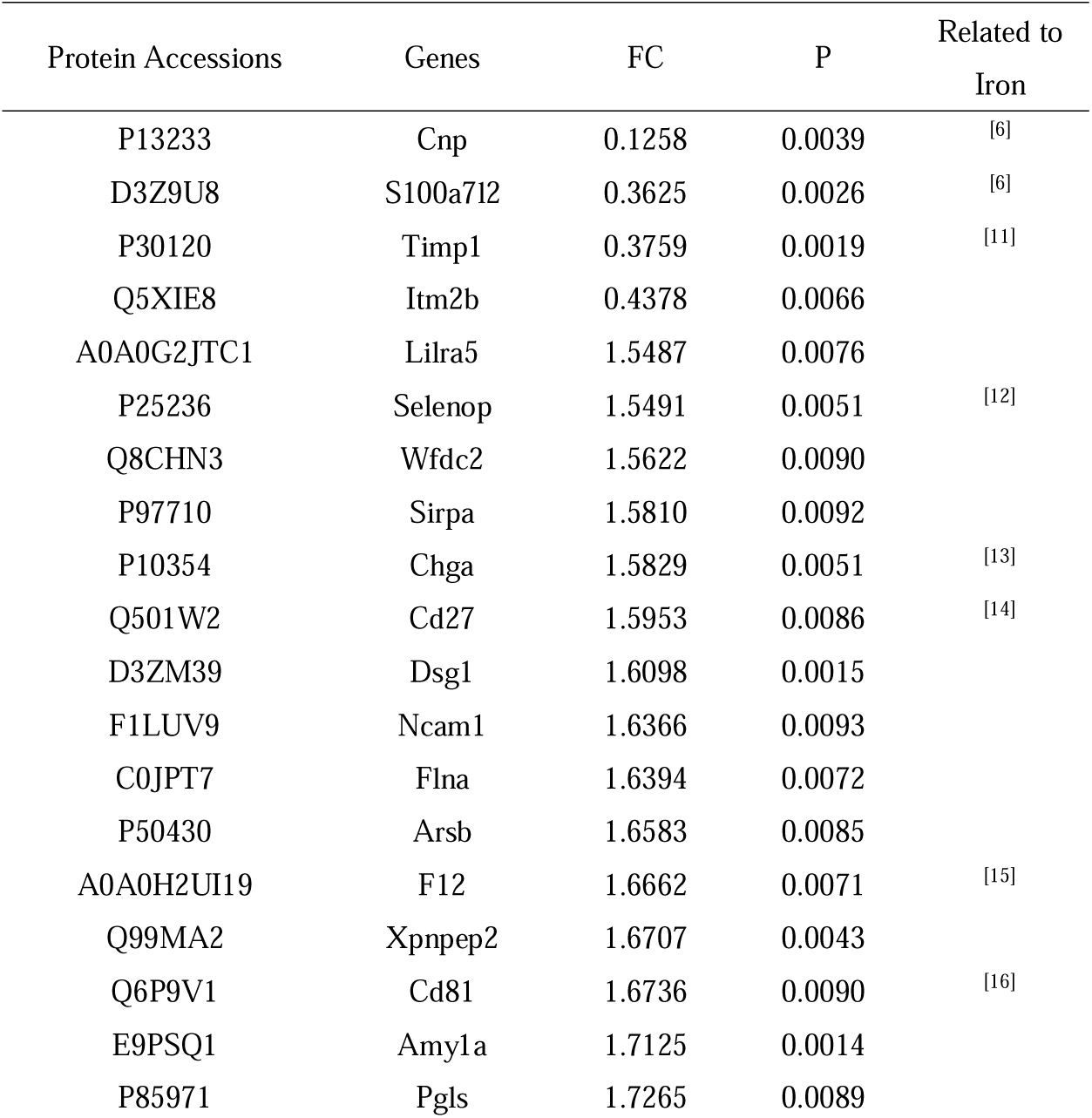
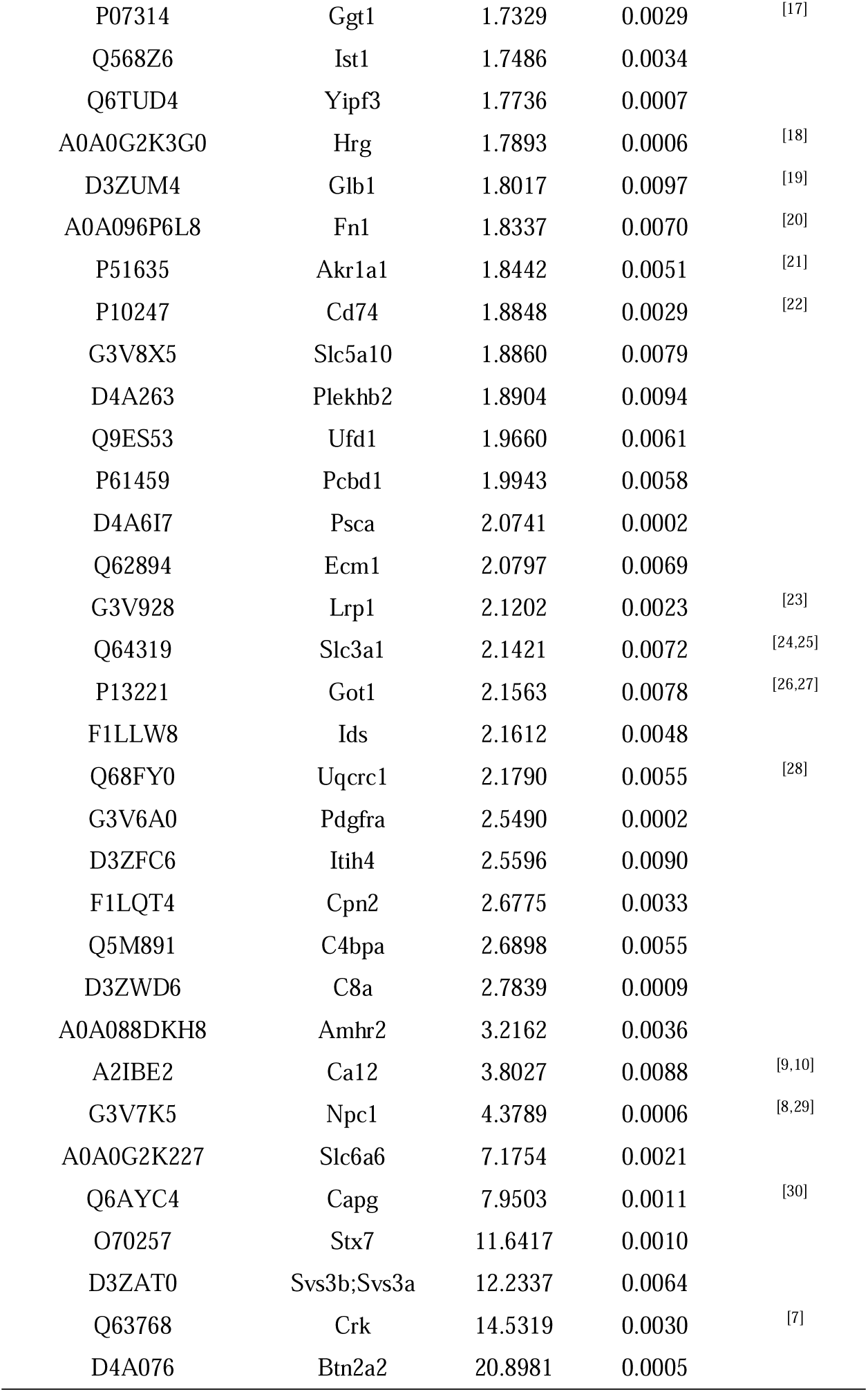
Differential proteins with significant changes in the comparative analysis of the Fe-D0 and Fe-D4 groups (p-value < 0.01, FC > 1.5 or < 0.67)

The 52 differential proteins were analyzed for protein function and literature searched using the PubMed database, and the literature showing the correlation of the differential proteins with iron is listed in the table.

After ingestion of polysaccharide iron, the down-regulated proteins in the urine included 2’,3’-cyclic-nucleotide 3’-phosphodiesterase (CNPase), S100 calcium binding protein A7 like 2 (S100A7l2), Tissue inhibitor of metalloproteinases 1 (TIMP-1), Integral membrane protein 2B. CNPase (FC=0.13) is a myelin marker. S100A7 is a protein capable of inducing immunomodulatory activity. It was found that the expression levels of CNPase and S100 calcium-binding protein were reduced in the offspring brains of rats on an iron-deficient diet during pregnancy, indicating that the availability of iron affects the development of oligodendrocytes^[6]^. TIMP-1 was overexpressed in rats treated with the iron-chelating agent Deferiprone compared to control rats.

The FC of the proto-oncogene c-Crk articulation molecule (p38) was 14.53. p38 protein expression was up-regulated in iron-overloaded bone marrow mesenchymal stem cells (BMSC)^[7]^. The FC of NPC intracellular cholesterol transporter 1 was 4.38. It was demonstrated that iron overload increased intracellular cholesterol^[8]^. The FC of Carbonic anhydrase (CA) is 3.8. Studies in experimental animals have shown that elevated oxidative stress in erythrocytes leads to the formation of autoantibodies against carbonic anhydrase and anemia^[9]^. Carbonic anhydrase may have an interfering effect on iron metabolism^[10]^.

#### 3.3.2 Randomized grouping results

To determine the likelihood of random generation of differential proteins identified by group comparisons, we validated the random grouping of total proteins identified from 10 samples from both groups, applying the same criteria for screening differential proteins: FC ≥ 1.5 or ≤ 0.67, P < 0.01, and performing 126 random groupings yielded an average of 10.82 differential proteins, with a proportion of randomly identified proteins of 21.15%, indicating that at least 79.85% proportion of the differential proteins were not due to randomness. The results of the randomized grouping test are shown in Table 2, and the 52 differential proteins (FC ≥ 1.5 or ≤ 0.67, P < 0.01) obtained from our screening had a low probability of being randomly generated, and the results showed that these differential proteins were indeed associated with short-term intake of polysaccharide iron complex supplements.

**Table 2.**
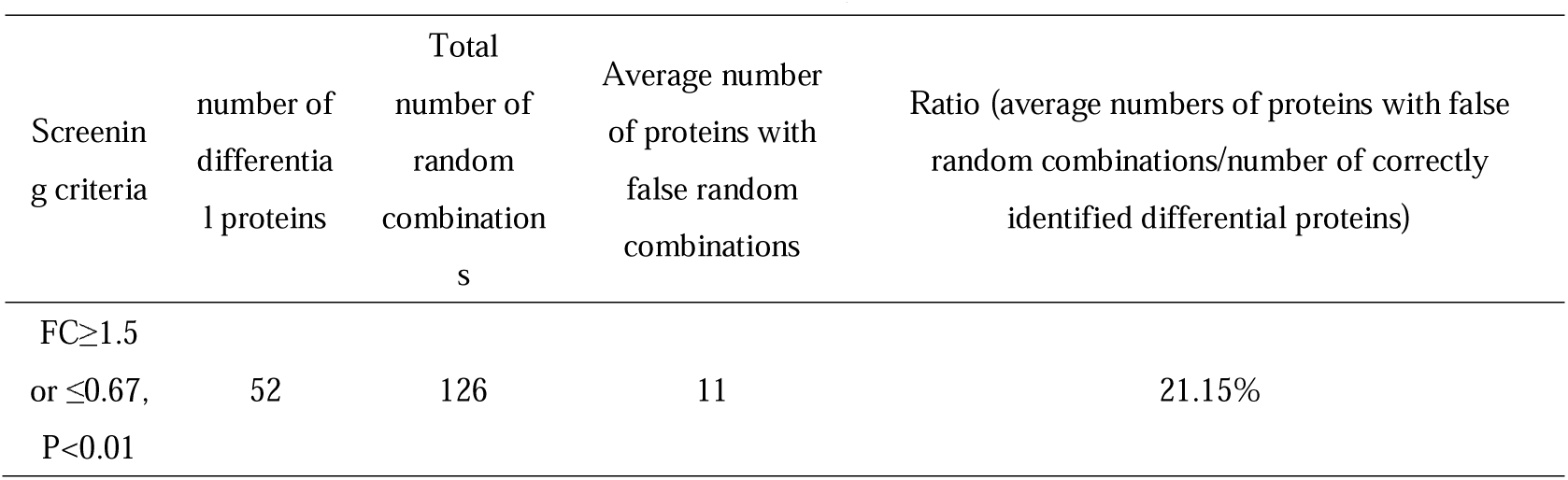
Results of randomized grouping of Fe-D0 and Fe-D4 groups according to the screening conditions of FC ≥ 1.5 or ≤ 0.67, P < 0.01.

| Screening criteria | number of differential proteins | Total number of random combinations | Average number of proteins with false random combinations | Ratio (average numbers of proteins with false random combinations/number of correctly identified differential proteins) |
| --- | --- | --- | --- | --- |
| $FC \geq 1.5$ or $\leq 0.67$, $P < 0.01$ | 52 | 126 | 11 | 21.15% |

We also verified the random grouping of total proteins identified from 10 samples in both groups according to the screening conditions of FC ≥ 1.5 or ≤ 0.67, P < 0.05. The average number of differential proteins obtained from 126 random groupings performed was 55, and the percentage of randomly identified proteins was 35.08%, indicating that at least 65% of the percentage of differential proteins were not due to randomness. The 157 differential proteins (FC ≥ 1.5 or ≤ 0.67, P < 0.05) obtained from our screening had a low probability of being randomly generated.

#### 3.3.3 Biological pathway analysis

The 157 differential proteins (P-value < 0.05, FC > 1.5 or < 0.67) were imported into the DAVID database and enriched to 53 biological processes (BPs), as shown in Table 3.

**Table 3.** Biological process (BP) enrichment analysis of differential proteins (P-value < 0.05, FC > 1.5 or < 0.67) in the Fe-D0 and Fe-D4 groups (P-value < 0.05, ordered by P-value from smallest to largest)

| Term | Count | % | P-Value | Related to Iron |
| --- | --- | --- | --- | --- |
| complement activation | 5 | 3.4 | 0.000025 | [47] |
| aging | 12 | 8.1 | 0.000041 | [32] |
| glucose transmembrane transport | 4 | 2.7 | 0.00095 | [33] |
| complement activation, classical pathway | 5 | 3.4 | 0.0013 | [47] |
| response to estrogen | 6 | 4.1 | 0.0016 | [34,35] |
| positive regulation of peptidyl-tyrosine phosphorylation | 6 | 4.1 | 0.0018 |  |
| metanephric proximal tubule development | 3 | 2 | 0.0022 | [48] |
| regulation of cell shape | 6 | 4.1 | 0.0044 |  |
| establishment of endothelial barrier | 3 | 2 | 0.011 | [37] |
| organic anion transport | 3 | 2 | 0.011 |  |
| negative regulation of apoptotic process | 11 | 7.4 | 0.014 | [38] |
| transmembrane transport | 7 | 4.7 | 0.014 |  |
| protein stabilization | 6 | 4.1 | 0.015 |  |
| response to iron ion | 3 | 2 | 0.016 |  |
| carbohydrate metabolic process | 5 | 3.4 | 0.017 | [49] |
| Cellular response to dexamethasone stimulus | 4 | 2.7 | 0.018 | [36] |
| positive regulation of fibroblast proliferation | 4 | 2.7 | 0.021 | [50] |
| negative regulation of intestinal absorption | 2 | 1.4 | 0.022 |  |
| cellular carbohydrate metabolic process | 2 | 1.4 | 0.022 | [49] |
| urate salt excretion | 2 | 1.4 | 0.022 | [51] |
| camera-type eye development | 4 | 2.7 | 0.023 | [52] |
| neuron migration | 5 | 3.4 | 0.026 | [53] |
| response to ethanol | 6 | 4.1 | 0.026 | [54,55] |
| zymogen activation | 3 | 2 | 0.026 | [56] |
| cell adhesion mediated by integrin | 3 | 2 | 0.028 | [50,57] |
| Defense response to Gram-negative bacterium | 4 | 2.7 | 0.028 | [58] |
| T cell activation via T cell receptor contact with antigen bound to MHC molecule on antigen presenting cell | 2 | 1.4 | 0.029 |  |
| transepithelial water transport | 2 | 1.4 | 0.029 |  |
| regulation of macrophage migration | 2 | 1.4 | 0.029 | [59] |
| glucuronate catabolic process to xylulose 5-phosphate | 2 | 1.4 | 0.029 | [33] |
| glycosylceramide catabolic process | 2 | 1.4 | 0.029 | [60] |
| Cellular response to interleukin-6 | 3 | 2 | 0.029 | [39,40] |
| cell-matrix adhesion | 4 | 2.7 | 0.03 | [50,57] |
| sodium ion transport | 4 | 2.7 | 0.031 | [41] |
| membrane fusion | 3 | 2 | 0.031 |  |
| Antimicrobial humoral immune response mediated by antimicrobial peptide | 4 | 2.7 | 0.033 |  |
| acute-phase response | 3 | 2 | 0.034 |  |
| dermatan sulfate catabolic process | 2 | 1.4 | 0.036 | [61] |
| L-ascorbic acid biosynthetic process | 2 | 1.4 | 0.036 | [42] |
| protein localization to plasma membrane | 5 | 3.4 | 0.036 |  |
| regulation of actin cytoskeleton organization | 4 | 2.7 | 3.60E-02 |  |
| killing of cells of other organism | 3 | 2 | 3.70E-02 |  |
| innate immune response | 8 | 5.4 | 3.70E-02 | [43] |
| hematopoietic progenitor cell differentiation | 4 | 2.7 | 4.20E-02 | [44] |
| positive regulation of substrate adhesion-dependent cell spreading | 3 | 2 | 4.20E-02 | [50,57] |
| Cellular response to inorganic substance | 3 | 2 | 4.20E-02 |  |
| ossification | 4 | 2.7 | 4.30E-02 | [45] |
| L-cystine transport | 2 | 1.4 | 4.30E-02 | [46] |
| glycoside catabolic process | 2 | 1.4 | 4.30E-02 |  |
| establishment of Sertoli cell barrier | 2 | 1.4 | 4.30E-02 |  |
| membrane raft organization | 2 | 1.4 | 4.30E-02 |  |
| response to inorganic substance | 3 | 2 | 4.40E-02 |  |
| Cellular response to transforming growth factor beta stimulus | 4 | 2.7 | 4.90E-02 | [62] |

Several biological pathways have been reported to be associated with the biological functions of iron. For example, complement activation, glucose transmembrane transport, response to estrogen, establishment of endothelial barrier, negative regulation of apoptotic processes, response to iron ions, carbohydrate metabolic processes, zymogen activation, cellular response to interleukin-6, sodium ion transport, cellular matrix adhesion, and hematopoietic progenitor cell differentiation.

According to the literature, intravenous iron administration induces complement activation in vivo^[31]^ . Dysregulation of iron metabolism affects aging^[32]^ . Systemic, cellular iron and glucose metabolism pathways are interrelated^[33]^ . Elevated estrogen levels are associated with increased systemically available iron^[34]^ . Estrogen administration upregulates transferrin^[35]^ . Reduced hepatic iron concentration in rats chronically administered dexamethasone^[36]^ . Intracellular iron chelation enhances endothelial barrier function^[37]^ . Iron induces reactive oxygen species (ROS) production and apoptosis^[38]^ . Lower serum iron levels are significantly associated with higher serum IL-6 levels, which promote iron death in bronchial epithelial cells by inducing ROS-dependent lipid peroxidation and disrupting iron homeostasis^[39,40]^ . Iron accumulation in bronchial epithelial cells is dependent on sodium transport^[41]^. L-ascorbic acid promotes iron uptake^[42]^ . L-ascorbic acid promotes iron absorption. Host antimicrobial mechanisms reduce iron availability to pathogens. there are various ferritins that influence the innate immune response^[43]^ . Ferritin has multiple effects on human hematopoietic progenitor cells. Ferritin inhibits the growth of human hematopoietic progenitor cells in vitro and the proliferation of T lymphocytes in vitro.^[44]^ . Iron overload inhibits endochondral ossification^[45]^ . Iron regulates L-cystine uptake and downstream GSH production in two types of mammalian cells^[46]^ . Due to space limitations, other biological processes and their relevance to iron are listed in the table.

#### 3.3.4 Molecular function and KEGG pathway analysis

The 157 differential proteins (P-value <0.05, FC>1.5 or <0.67) were imported into the DAVID database and enriched to 23 molecular functions, as shown in Table 4.

**Table 4.**
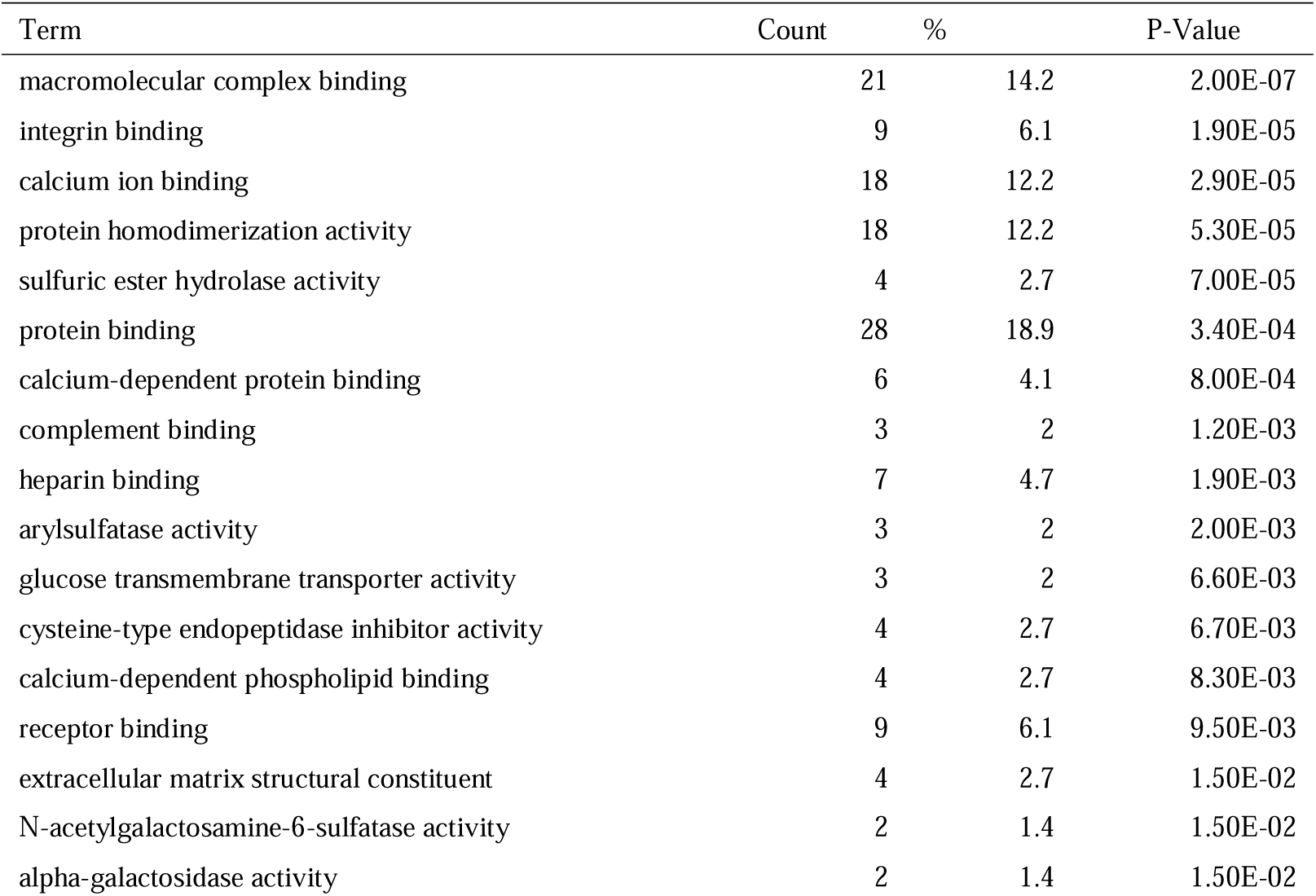
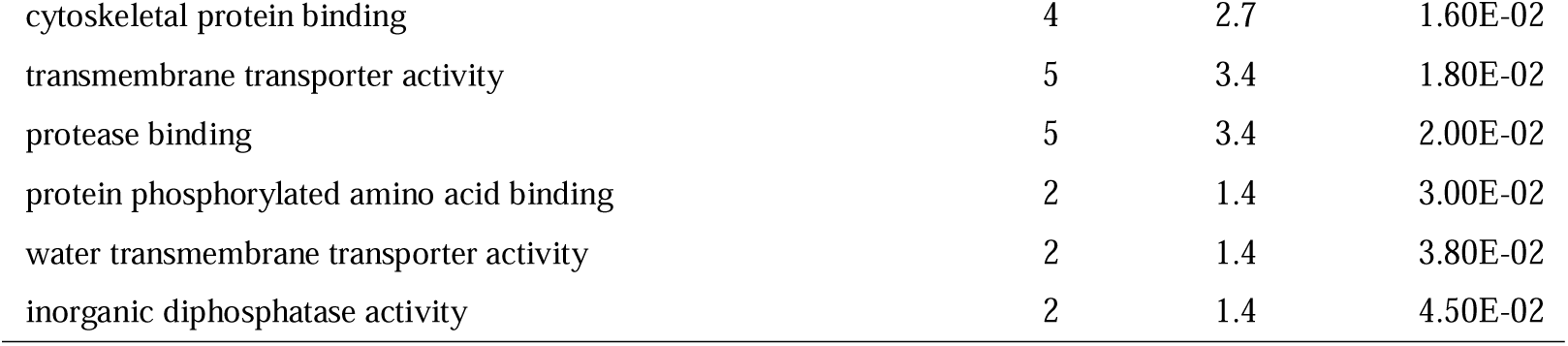
Molecular function (MF) enrichment analysis of differential proteins (P-value < 0.05, FC > 1.5 or < 0.67) in the Fe-D0 and Fe-D4 groups (P-value < 0.05, ordered by P-value from smallest to largest)

| Term | Count | % | P-Value |
| --- | --- | --- | --- |
| macromolecular complex binding | 21 | 14.2 | 2.00E-07 |
| integrin binding | 9 | 6.1 | 1.90E-05 |
| calcium ion binding | 18 | 12.2 | 2.90E-05 |
| protein homodimerization activity | 18 | 12.2 | 5.30E-05 |
| sulfuric ester hydrolase activity | 4 | 2.7 | 7.00E-05 |
| protein binding | 28 | 18.9 | 3.40E-04 |
| calcium-dependent protein binding | 6 | 4.1 | 8.00E-04 |
| complement binding | 3 | 2 | 1.20E-03 |
| heparin binding | 7 | 4.7 | 1.90E-03 |
| arylsulfatase activity | 3 | 2 | 2.00E-03 |
| glucose transmembrane transporter activity | 3 | 2 | 6.60E-03 |
| cysteine-type endopeptidase inhibitor activity | 4 | 2.7 | 6.70E-03 |
| calcium-dependent phospholipid binding | 4 | 2.7 | 8.30E-03 |
| receptor binding | 9 | 6.1 | 9.50E-03 |
| extracellular matrix structural constituent | 4 | 2.7 | 1.50E-02 |
| N-acetylgalactosamine-6-sulfatase activity | 2 | 1.4 | 1.50E-02 |
| alpha-galactosidase activity | 2 | 1.4 | 1.50E-02 |
| cytoskeletal protein binding | 4 | 2.7 | 1.60E-02 |
| transmembrane transporter activity | 5 | 3.4 | 1.80E-02 |
| protease binding | 5 | 3.4 | 2.00E-02 |
| protein phosphorylated amino acid binding | 2 | 1.4 | 3.00E-02 |
| water transmembrane transporter activity | 2 | 1.4 | 3.80E-02 |
| inorganic diphosphatase activity | 2 | 1.4 | 4.50E-02 |

157 differential proteins (P value <0.05, FC >1.5 or <0.67) were imported into the DAVID database and enriched to 10 KEGG pathways. KEGG pathways enriched to include lysosome, complement and coagulation cascades, focal adhesion, glycosaminoglycan degradation, regulation of the actin cytoskeleton, amebiasis, malaria, leukocyte transendothelial migration, systemic lupus erythematosus, and metabolic pathways. (Table 5)

**Table 5.**
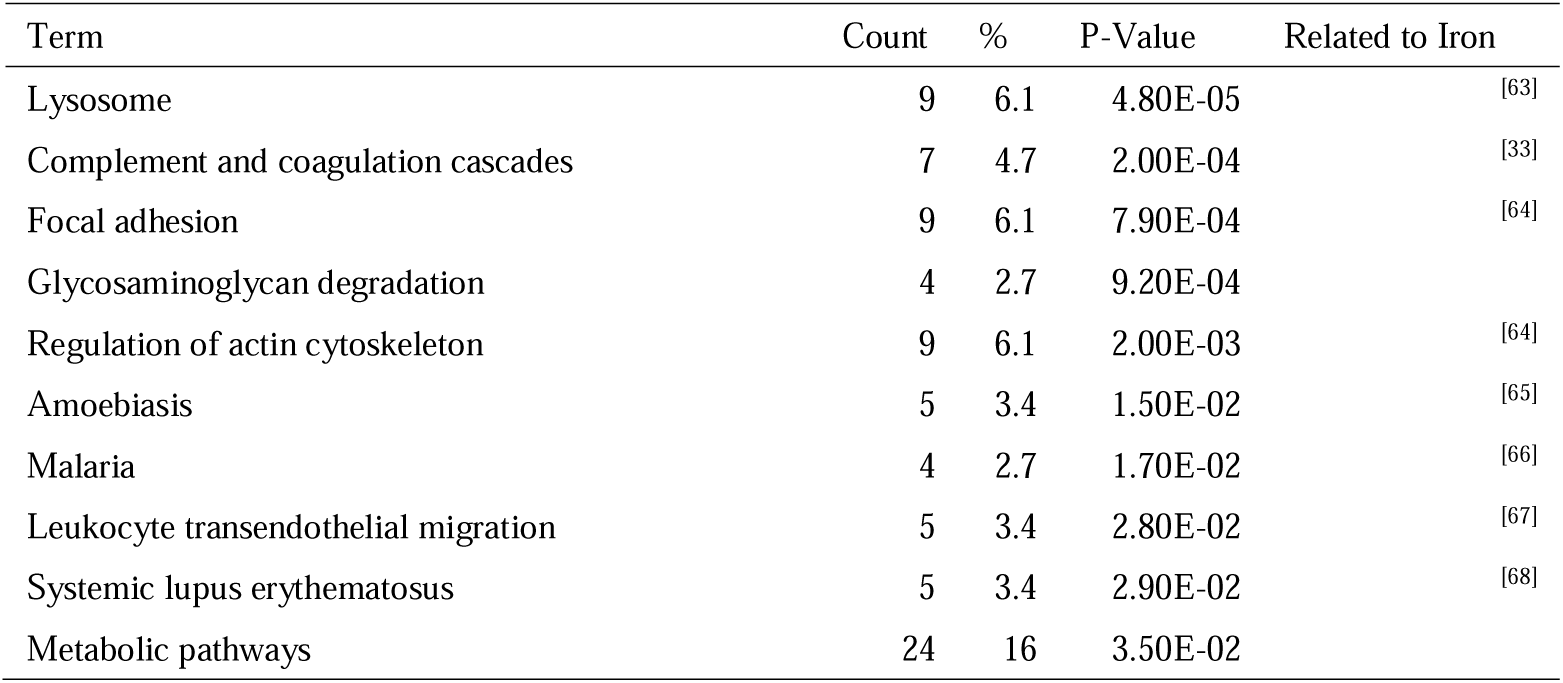
KEGG pathway enrichment analysis of differential proteins (P-value < 0.05, FC > 1.5 or < 0.67) in Fe-D0 and Fe-D4 groups (P-value < 0.05, ordered by P-value from smallest to largest)

Lysosomes are the main regulators of iron metabolism^[63]^ . Intravenous iron preparations induce complement activation in vivo^[33]^ . High intracellular iron oxide nanoparticle concentration affects cytoskeletal and adhesion patch kinase-mediated signaling^[64]^ . Administration of iron greatly increases susceptibility to amoebiasis in pastoralists with dietary iron deficiency^[65]^ . Iron is a cofactor in Plasmodium falciparum development^[66]^ . Iron sucrose and iron gluconate significantly inhibited transendothelial migration of polymorphonuclear leukocytes (PMN)^[67]^ . Many studies have demonstrated the important role of iron in the immune response, and there is growing evidence that iron homeostasis may be aberrant in the chronic inflammatory state of systemic lupus erythematosus (SLE)^[68]^ .

### 3.4 Individual comparison

#### 3.4.1 Screening for Differential Proteins

The urine proteome is very sensitive to changes in the state of the organism and can also be influenced to some extent by genetic factors^[69]^, age^[70–72]^, gender^[73,74]^, ethnicity^[75]^, region^[76]^ , exercise^[77,78]^, dietary habits, mental status, circadian rhythms, and medication use^[79,80]^, and other environmental factors that exhibit variability within the same individual and among individuals^[81,82]^. Animal models are easy to control variables. Animal models are easy to control variables and can reduce variation due to extraneous variables in human urine samples, but some variation exists even between individuals of the same species. Therefore, this study used its own before-and-after control analysis method, which can reduce the effect of individual variability and help to identify latent important information.

Urine proteomes before-after gavage of each rat were compared individually. Specific analyses for self before and after comparisons were performed as follows: missing values were replaced with 0. Three-needle replicates of the pre-gavage sample (D0) were compared two-tailed and pairwise with three-needle replicates of the sample on day 4 of gavage (D4) for each rat, and the screening for differential proteins conditioned on a P-value of <0.05 for t-test analysis, a fold change Fold change (FC) >1.5 or < 0.67.

The screening results were as follows: 194 differential proteins were screened in rat #1, 368 differential proteins in rat #2, 520 differential proteins in rat #3, 230 differential proteins in rat #4, and 148 differential proteins in rat #5.

#### 3.4.2 Shared biological processes, molecular functions, pathway analysis

Differential proteins from each of the five rats were functionally annotated using the DAVID database with a screening condition of p<0.05. The overlap of biological processes, molecular functions, and pathways of the five rats was also analyzed using a Wayne diagram.

Rat #1 was enriched to 126 biological processes; rat #2 was enriched to 163 biological processes; rat #3 was enriched to 212 biological processes; rat #4 was enriched to 167 biological processes; and rat #5 was enriched to 77 biological processes.

There were three biological processes shared among five rats (100% of the total experimental group), including carbohydrate metabolic processes, aging, and cell-matrix adhesion. The literature shows that dysregulation of iron metabolism and aging^[32]^ , glucose metabolism pathways are interrelated^[33]^ and that iron death is associated with multiple signaling pathways such as cell adhesion^[50,57]^ .

Sixteen biological processes were common to four rats (80% of the total experimental group). These biological processes are shown in a table along with the literature related to iron. In addition, biological processes such as the response to iron ions and the regulation of eIF2 α phosphorylation by hemoglobin were shared in 3 rats (60% of the total experimental group).

Rat #1 was enriched to 35 molecular functions; rat #2 was enriched to 67 molecular functions; rat #3 was enriched to 75 molecular functions; rat #4 was enriched to 61 molecular functions; and rat #5 was enriched to 31 molecular functions.

Three molecular functions were shared in five rats (100% of the total experimental group), including protein binding, macromolecular complex binding, and calcium ion binding. 11 molecular functions were shared in four rats (80% of the total experimental group), including hemoglobin beta binding. Iron is an important component of hemoglobin synthesis in vivo and is essential for oxygen transport and cellular respiration.

Pathway enrichment analysis was performed using the Kyoto Encyclopedia of genes and genomes database ( Kyoto encyclopedia of genes and genomes, KEGG ) using the DAVID website. 30 KEGG pathways were enriched in rat #1; 41 KEGG pathways were enriched in rat #2; 49 KEGG pathways in rat #3; 10 KEGG pathways were enriched in rat #4; and 35 KEGG pathways in rat #5. enriched to 35 KEGG pathways; rat #5 enriched to 10 KEGG pathways.

KEGG pathways that were common to 5 rats (100% of the total experimental group) were lysosomes, phagosomes. 5 KEGG pathways were common to 4 rats (80% of the total experimental group) including malaria, endocytosis, African trypanosomiasis, Staphylococcus aureus infection, sphingolipid metabolism. Literature suggesting pathway association with iron is tabulated.

#### 3.4.3 Analysis of differential proteins jointly up- or down-regulated in multiple rats

Differential proteins obtained from the before-and-after comparison of each rat were divided into up-regulated and down-regulated according to FC. Compared with the pre-gastric sample (D0), 129, 309, 425, 148, 69 up-regulated differential proteins were identified in the post-gastric sample (D4) in 5 rats (FC>1.5, P<0.05), and 65, 59, 95, 82, 79 down-regulated differential proteins were identified in the post-gastric sample in 5 rats (FC<0.67, P<0.05). The overlap of differential proteins identified before and after gavage in five rats was demonstrated using Wayne diagrams, as shown in Figures 4 and 5. Differential protein names and overlaps are listed in the Supplementary Table.

**Figure 4.**
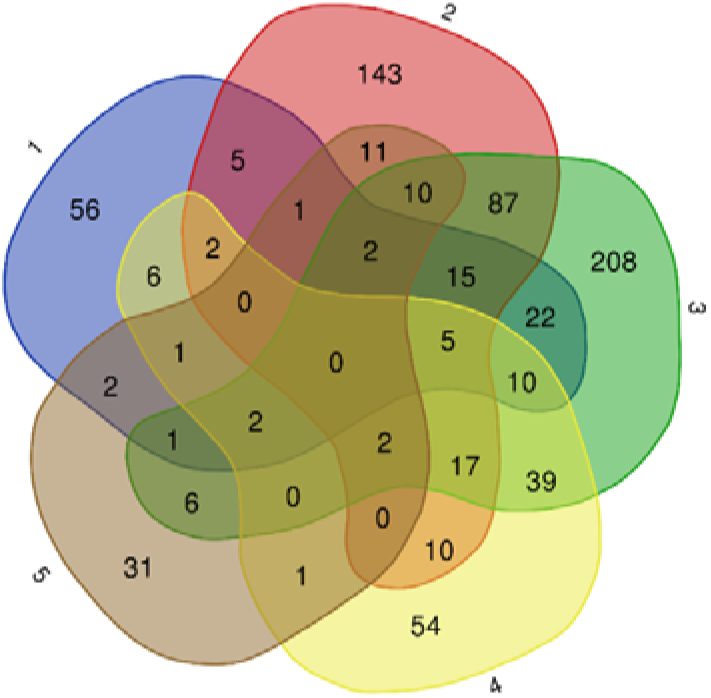
Wayne plots of up-regulated differential proteins produced by 5 rats before and after their own control (FC>1.5, P<0.05)

**Figure 5.**
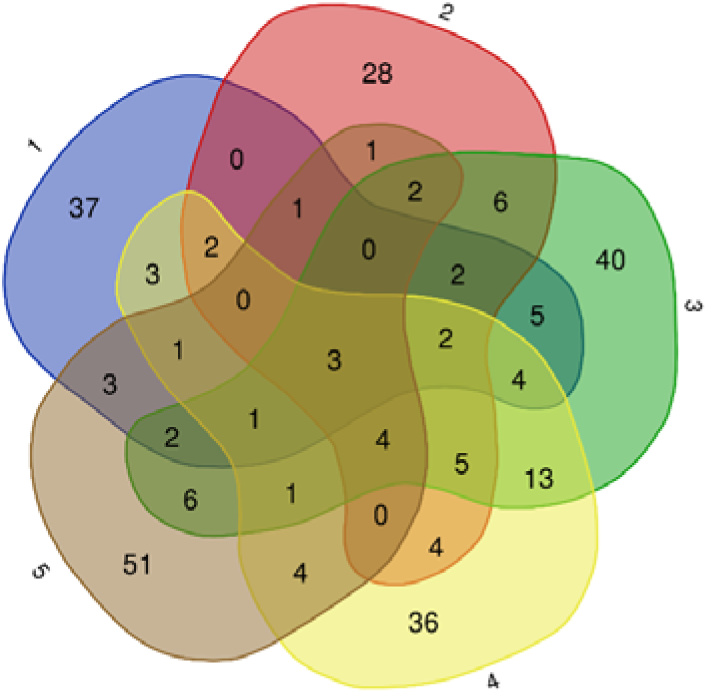
Down-regulated differential proteins produced by the control before and after the 5 rats themselves (FC<0.67,P<0.05) Wayne plots

Detailed searches and analyses were performed for those of these differential proteins that were jointly up- or down-regulated in four or five rats, as shown in Table 9.

**Table 6.** Biological processes (BP) shared in 4 or 5 rats (DAVID database GO analysis) Rats Biological Process (BP)

| Rats | Biological Process (BP) | Related to Iron |
| --- | --- | --- |
| 1 2 3 4 5 | carbohydrate metabolic process | [33] |
| 1 2 3 4 5 | aging | [32] |
| 1 2 3 4 5 | cell-matrix adhesion | [83] |
| 1 2 3 4 | negative regulation of cysteine-type endopeptidase activity | [46] |
| 1 2 3 4 | negative regulation of endopeptidase activity | [56] |
| 1 2 3 4 | positive regulation of fibroblast proliferation | [50] |
| 1 2 3 4 | lipid metabolic process | [84] |
| 1 2 3 4 | cell adhesion mediated by integrin | [50,57] |
| 1 2 3 4 | response to estrogen | [34,35] |
| 1 2 3 4 | glomerular filtration | [85,86] |
| 1 2 3 5 | phagocytosis, engulfment | [87] |
| 1 3 4 5 | positive regulation of cell migration | [88] |
| 1 3 4 5 | positive regulation of ERK1 and ERK2 cascade | [89] |
| 1 3 4 5 | Cellular response to lipopolysaccharide | [90,91] |
| 1 3 4 5 | positive regulation of protein kinase B signaling | [92] |
| 2 3 4 5 | zymogen activation | [56] |
| 2 3 4 5 | regulation of systemic arterial blood pressure | [93] |
| 2 3 4 5 | complement activation, classical pathway | [47] |
| 2 3 4 5 | glutathione metabolic process | [94] |

**Table 7.** Molecular functions (MF) shared in 4 or 5 rats (DAVID database GO analysis) Rats Molecular function (MF)

| Rats | Molecular function (MF) |
| --- | --- |
| 1 2 3 4 5 | protein binding |
| 1 2 3 4 5 | macromolecular complex binding |
| 1 2 3 4 5 | calcium ion binding |
| 1 2 3 4 | proton-transporting ATPase activity, rotational mechanism |
| 1 2 3 4 | hydrolase activity |
| 1 2 3 4 | protease binding |
| 1 2 3 4 | cysteine-type endopeptidase inhibitor activity |
| 1 2 3 4 | serine-type endopeptidase inhibitor activity |
| 1 2 3 4 | phosphatidylserine binding |
| 1 2 4 5 | hemoglobin beta binding |
| 1 3 4 5 | endopeptidase inhibitor activity |
| 2 3 4 5 | peptidase activity |
| 2 3 4 5 | receptor binding |
| 2 3 4 5 | serine-type endopeptidase activity |

**Table 8.** KEGG pathways shared in 4 or 5 rats (DAVID database GO analysis)

| Rats | Kyoto encyclopedia of genes and genomes (KEGG) pathway | Related to Iron |
| --- | --- | --- |
| 1 2 3 4 5 | Lysosome | [63] |
| 1 2 3 4 5 | Phagosome | [95] |
| 1 2 4 5 | Malaria | [66] |
| 1 2 4 5 | Endocytosis | [96] |
| 1 3 4 5 | African trypanosomiasis | [97] |
| 1 3 4 5 | Staphylococcus aureus infection | [58] |
| 2 3 4 5 | Sphingolipid metabolism | [98] |

**Table 9.**
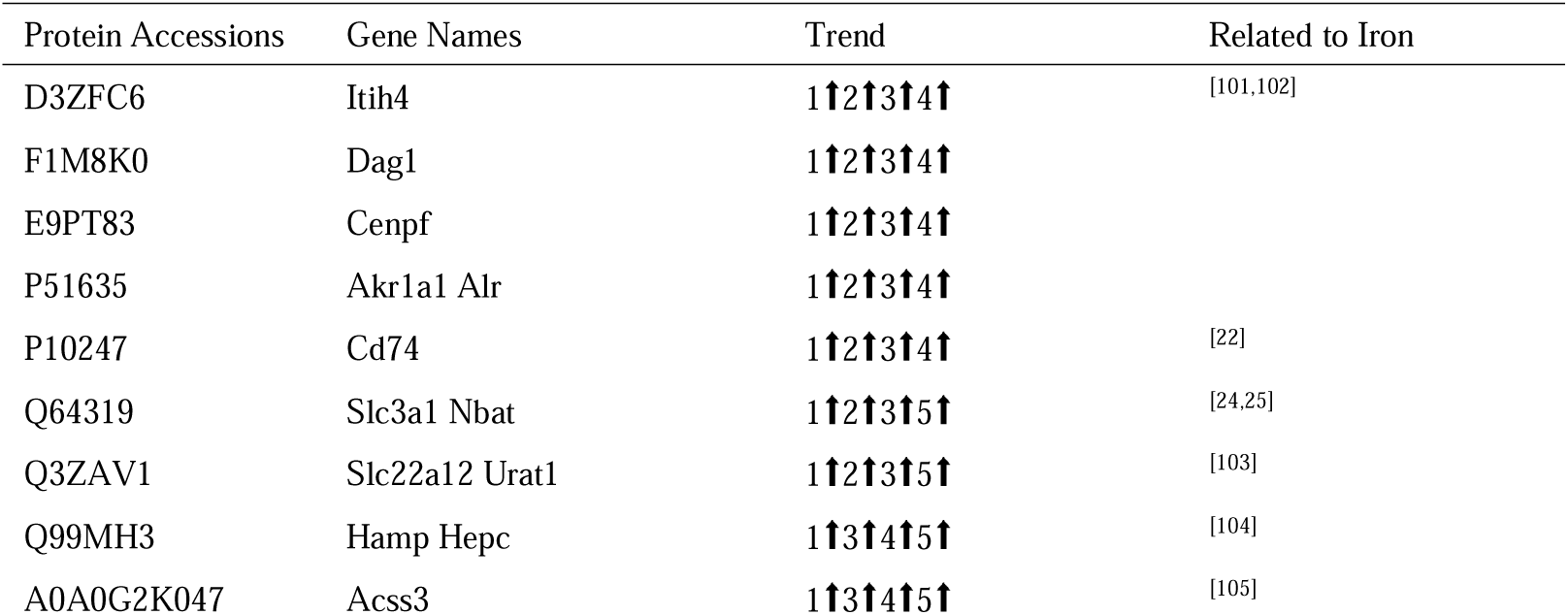
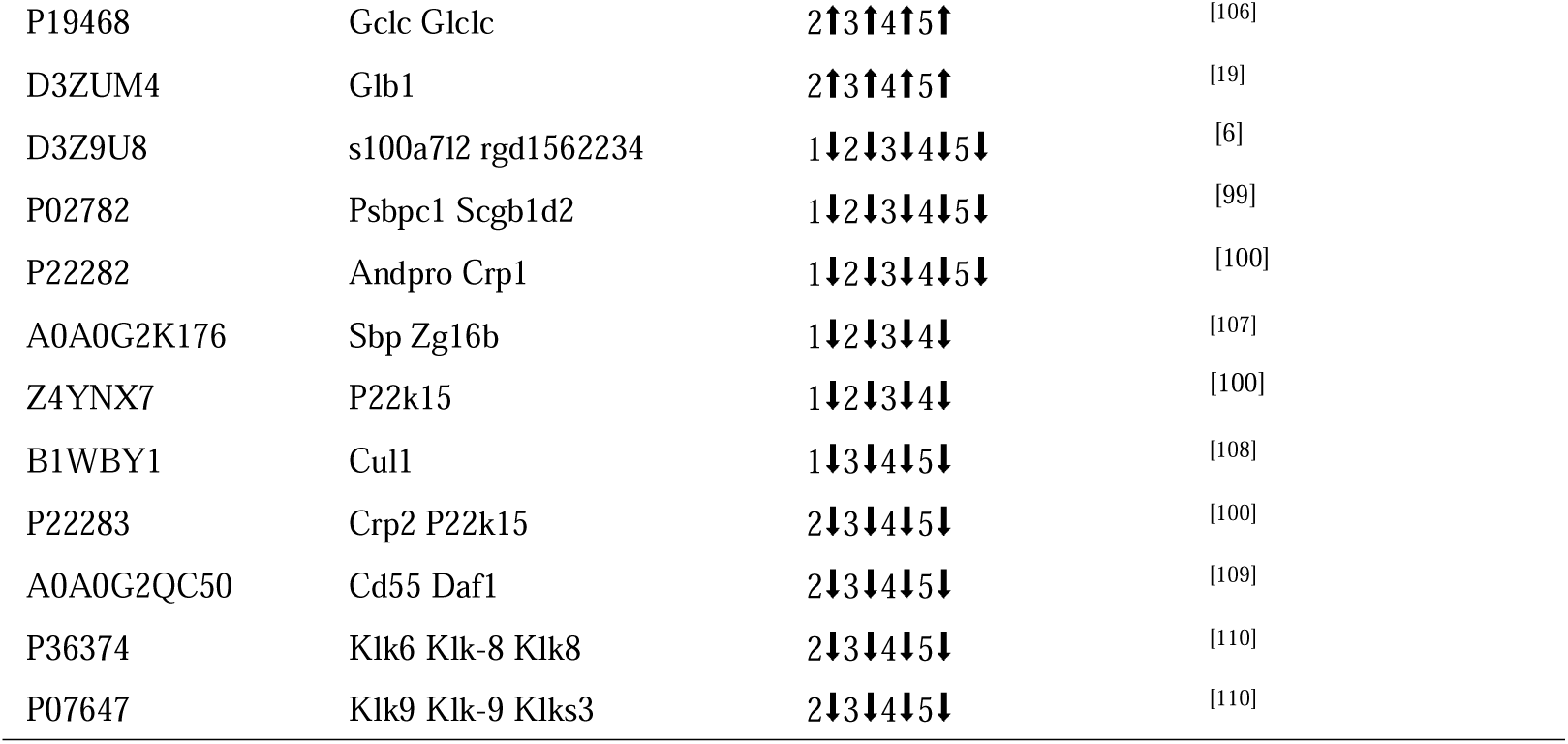
Differential proteins jointly up- or down-regulated in 4 or 5 rats.

S100 calcium binding protein A7 like 2 (S100A7l2), prostatic steroid-binding protein C1, and cystatin-related protein 1 were jointly down-regulated in five rats. Expression of S100 calcium-binding protein was decreased in the brains of offspring from rats fed an iron-deficient diet during pregnancy^[6]^. Prostate epithelial cells synthesize hepcidin, and hepcidin synthesis and secretion were significantly increased in prostate cancer cells and tissues^[99]^. Cystatin C is positively correlated with serum ferritin^[100]^. Cystatin C is positively correlated with serum ferritin. Seven proteins were down-regulated in four rats, including spermine binding protein, cystatin-related protein 2, Cullin 1, decay accelerating factor 1, prostatic glandular kallikrein-6, and submandibular glandular kallikrein-9. A review of the literature revealed that a variety of proteins (or families thereof) are associated with iron metabolism or ferritin, as detailed in Table 9. Eleven proteins were up-regulated in four rats, including hepcidin, which acts as a signaling molecule involved in the maintenance of iron homeostasis. H-2 class II histocompatibility antigen gamma chain, Neutral and basic amino acid transport protein rBAT, Solute carrier family 22 member 12, Acyl-CoA synthetase short-chain family member 3, Glutamate-cysteine ligase catalytic subunit, Beta-galactosidase and many other proteins were also searched for their association with iron metabolism or ferritin.

#### 3.4.4 Functional analysis of enrichment of differentially differentiated proteins jointly up- or down-regulated in multiple rats

Functional annotation was performed for differential proteins that were co-up- or down-regulated in 3, 4, or 5 rats, and for biological processes (Table 10), molecular functions (Fig. 6), and KEGG pathways (Table 11) that were enriched for these differential proteins.

**Figure 6.**
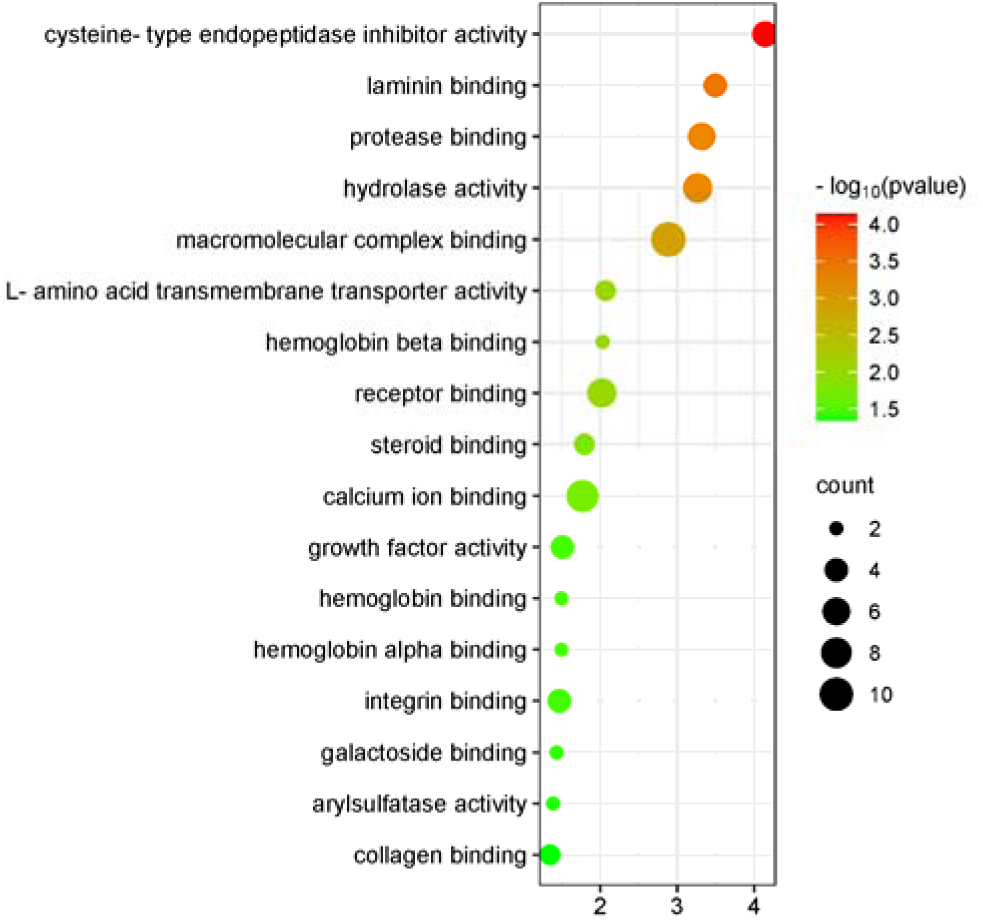
Molecular function (MF) enrichment analysis of proteins that were co-up- or down-regulated in 3 or more rats (DAVID database GO analysis)

**Table 10.**
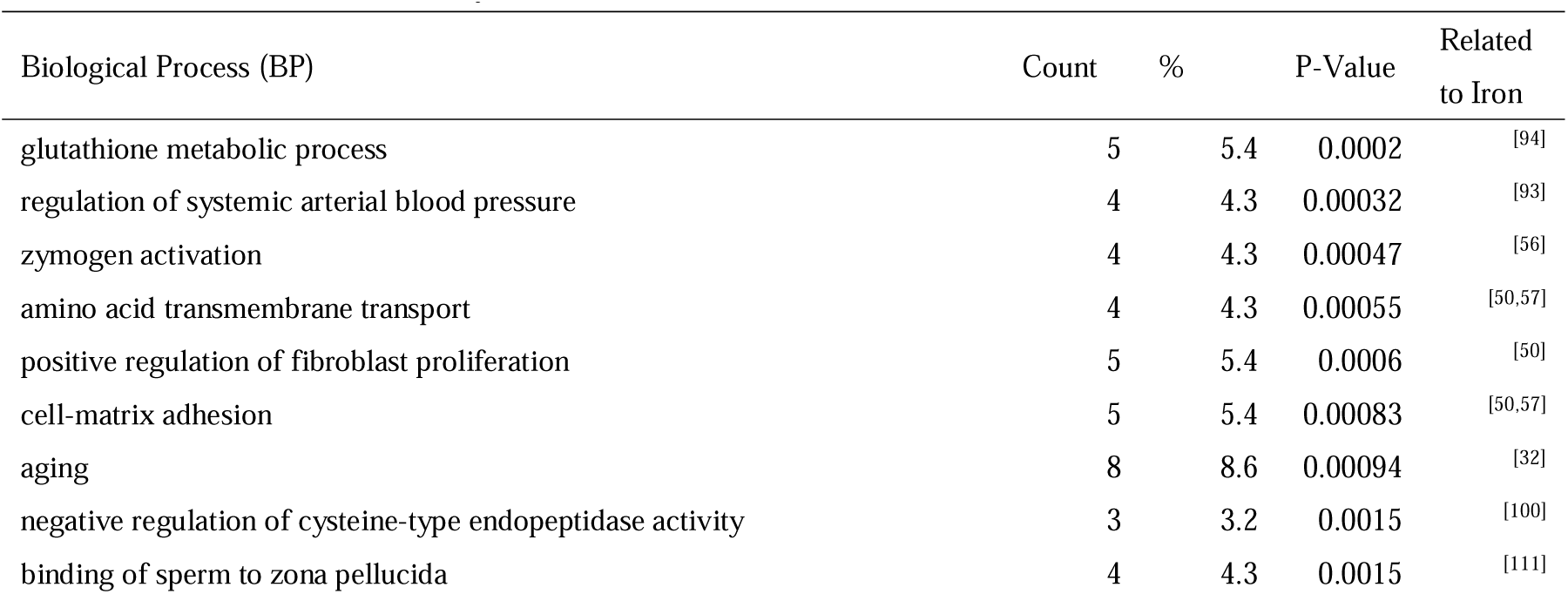
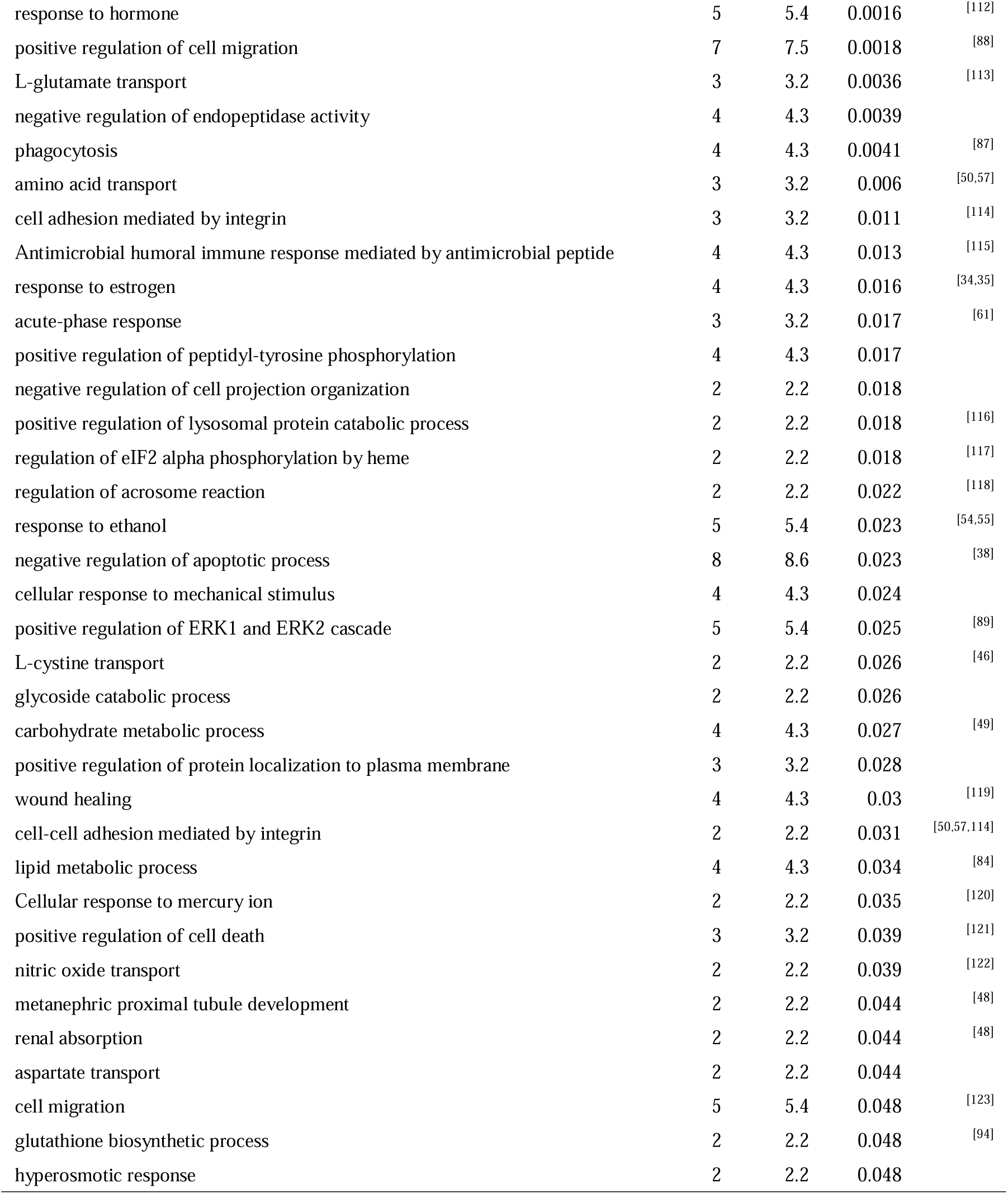
Biological process (BP) enrichment analysis of proteins commonly up- or down-regulated in 3 or more rats (DAVID database GO analysis)

**Table 11.** KEGG pathway enrichment analysis of proteins commonly up- or down-regulated in 3 or more rats (DAVID database GO analysis)

| KEGG Pathway | Count | % | P-Value | Related to Iron |
| --- | --- | --- | --- | --- |
| Lysosome | 7 | 7.5 | 0.000085 | [63] |
| Sphingolipid metabolism | 4 | 4.3 | 0.0035 | [98] |
| Other glycan degradation | 3 | 3.2 | 0.0043 | [124] |
| Glycosaminoglycan degradation | 3 | 3.2 | 0.0053 | [125] |
| Regulation of actin cytoskeleton | 6 | 6.5 | 0.0089 | [64] |
| Gap junction | 4 | 4.3 | 0.012 | [126] |
| Phagosome | 5 | 5.4 | 0.019 | [95] |
| African trypanosomiasis | 3 | 3.2 | 0.02 | [97] |
| Cholesterol metabolism | 3 | 3.2 | 0.031 | [84] |
| Malaria | 3 | 3.2 | 0.036 | [66] |

A total of 44 biological processes were enriched, and the correlation between the enriched biological processes and iron was searched, and the related literature is detailed in Table 10.

Fifteen of these biological processes overlapped with the results of the metagenomic analysis, including zymogen activation, positive regulation of fibroblast proliferation, cell matrix adhesion, aging, integrin-mediated cell adhesion, antimicrobial peptide-mediated antimicrobial humoral immune response, response to estrogens, acute-phase response, positive regulation of peptidyl-tyrosine phosphorylation, response to ethanol, negative regulation of apoptotic processes, L-cystine transport, glycosidic catabolic processes, carbohydrate metabolic processes, adrenal proximal tubular development.

In addition, biological processes such as glutathione metabolic processes, regulation of systemic arterial blood pressure, and regulation of eIF2α phosphorylation by heme were also enriched. Many biological processes are related to the biological functions of iron.

Seventeen molecular functions were enriched, including hemoglobin binding. Seven molecular functions, including cysteine-type endopeptidase inhibitor activity, protease binding, macromolecular complex binding, receptor binding, calcium ion binding, integrin binding, and aryl sulfatase activity, overlapped with the results of the group analysis.

Ten KEGG pathways were enriched, four of which overlapped with the results of the group-forming analysis, including lysosomes, glycosaminoglycan degradation, regulation of the actin cytoskeleton, and malaria. The KEGG pathways enriched for correlation with iron were searched, and the relevant literature is detailed in Table 11.

## 4 Perspective

Iron overload is usually defined as an excessive accumulation of iron in the body beyond normal metabolic needs. This condition may be caused by a variety of reasons, such as long-term iron supplementation, hereditary diseases (e.g., hereditary hemochromatosis), and chronic inflammatory states. In recent years, concerns have been raised regarding the occurrence of iron overload and its negative effects. Iron overload occurs all over the world, especially in the more economically developed regions, and has a serious impact on human (especially children’s) health and life safety. Iron overload affects lipid peroxidation, nutritional metabolism and is closely related to the development of cardiovascular diseases. The effects of iron overload on the health of living organisms are multifaceted and include, but are not limited to, increased intracellular oxidative stress, tissue damage, impaired organ function, which can lead to serious cardiovascular and neurological diseases.

In this study, rats were gavaged with polysaccharide iron complex at a dose of 28 mg/kg-d (by iron), which is equivalent to the dose used to prevent anemia in adults. According to literature research, the concentration of polysaccharide iron complex used in this study would require more than 4 weeks of gavage if applied to establish an iron overload model^[127]^. The present study was conducted in rats by gavage of polysaccharide iron complex (28 mg/kg-d iron) for 4 days with the aim of investigating the overall effect of short-term polysaccharide iron complex gavage on the organism. This study is expected to provide some clues for the prevention, diagnosis, treatment and monitoring of iron metabolism disorders (e.g., anemia due to iron deficiency and cardiovascular disease due to iron overload) and to fill the gap in the field of iron metabolism in the urine proteome.

In this study, two analytical methods were used: before-and-after comparison of itself and comparison of groups, which provided us with more comprehensive and reliable data validation. The application of the before-and-after comparison method reduces the influence of individual differences on the experimental results, improves the stability and reproducibility of the experiment, and is important for the credibility of the results. The mutual validation of the results obtained by the two analytical methods indicates that the urine proteome is able to reflect the effects of short-term intake of polysaccharide-iron complexes on the organism, making the results more credible.

The results of the study illustrate that the urine proteome of rats can show changes in iron-related proteins and biological functions after short-term intake of polysaccharide iron complexes. Short-term supplementation of polysaccharide iron complexes can affect the organism, and the urine proteome can reflect the overall changes of the organism in a comprehensive and systematic way. The present study provides clues for a deeper understanding of the metabolic process, mechanism of action, and biological function of iron in organisms from the perspective of urine proteomics, as well as new research perspectives and methodological insights for future related studies, which are potentially important for the prevention, diagnosis, treatment, and monitoring of iron metabolism disorder-related diseases.

